# From Space to Sequence and Back Again: Iterative DNA Proximity Ligation and its Applications to DNA-Based Imaging

**DOI:** 10.1101/470211

**Authors:** Alexander A. Boulgakov, Erhu Xiong, Sanchita Bhadra, Andrew D. Ellington, Edward M. Marcotte

## Abstract

We extend the concept of DNA proximity ligation from a single readout per oligonucleotide pair to multiple reversible, iterative ligations re-using the same oligonucleotide molecules. Using iterative proximity ligation (IPL), we can in principle capture multiple ligation events between each oligonucleotide and its various neighbors and thus recover a far richer knowledge about their relative positions than single, irreversible ligation events. IPL would thus act to sample and record local molecular neighborhoods. By integrating a unique DNA barcode into each participating oligonucleotide, we can catalog the individual ligation events and thus capture the positional information contained therein in a high throughput manner using next-generation DNA sequencing. We propose that by interpreting IPL sequencing results in the context of graph theory and by applying spring layout algorithms, we can recover geometric patterns of objects labeled by DNA. Using simulations, we demonstrate that we can in principle recover letter patterns photolithographed onto slide surfaces using only IPL sequencing data, illustrating how our technique maps complex spatial configurations into DNA sequences and then – using only this sequence information – recovers them. We complement our theoretical work with an experimental proof-of-concept of iterative proximity ligation on an oligonucleotide population.

## Introduction

Aside from its everyday task of carrying genetic information in organisms, DNA is also a wonderful molecule with broad utility for applications such as nanotechnology, computation, and information storage^1,2,3,4^. As our mastery of chemistries and enzymes to manipulate DNA has deepened, so has the breadth of its utility. The dominant development contributing to this trend has been the rapid growth in high-throughput DNA sequencing^5^, now providing us with a remarkable stream of new information about the world. DNA barcoding—the incorporation of specific, pre-designed DNA sequences into molecules as a means to identify them or copies of them across experiments—has also been a key innovation for accelerating the throughput of sequencing^6^ and as such sits at an interesting intersection of biology and information theory. One application of DNA barcoding is to label and thus track individual molecules as they are manipulated. This concept has been expanded to obtain information beyond tracking particle identities, for example to identifying their interaction partners^7^ and spatial positions^8^.

Proximity ligation assays^9^ are one of the technologies that leverage oligonucleotides, and our abilities to manipulate them, to detect spatial co-localization of molecules. A pair of proximity probes, usually antibodies, are conjugated to oligonucleotides and then incubated with the sample of interest. If two labeled probe targets are in sufficiently close proximity to each other, for example in the case of two proteins participating in the same complex, the pair of conjugated oligonucleotides are able to ligate and create a contiguous template that can then be amplified by (often circular) PCR, whose products can then in turn be visualized by fluorescent probes. Proximity ligation, however, is traditionally performed with one ligation step and thus captures only the pairwise proximity of individual probe pairs across the sample. Here, we extend the concept of proximity ligation to iterative rounds of reversible ligations on a population of barcoded DNA molecules, allowing us to capture information about the relative positions not only of paired probes in isolation but of the entire population as a whole.

Iterative proximity ligation (IPL) is based upon the observation that the processes of enzymatic ligation and digestion of DNA are ideally memoryless and hence can be applied repeatedly to the same molecule multiple times. We use this observation to consider multiple proximity ligations per oligonucleotide. Thus instead of being able to obtain only one piece of information, namely that two particular oligonucleotides are in sufficient proximity to each other to ligate, we can obtain many readouts of proximity information about an oligonucleotide and its neighbors, defining local molecular neighborhoods. These readouts can be interpreted as a graph, where barcoded oligonucleotides are nodes and observed ligations are edges (**Fig. 1**). The resulting IPL graph carries rich information about the labeled object(s). In particular, as we show below, applying spring layout algorithms to an IPL graph makes it possible to triangulate and determine the original position of each molecule, with positional errors on the order of the oligonucleotide chain lengths.

**Fig. 1.**
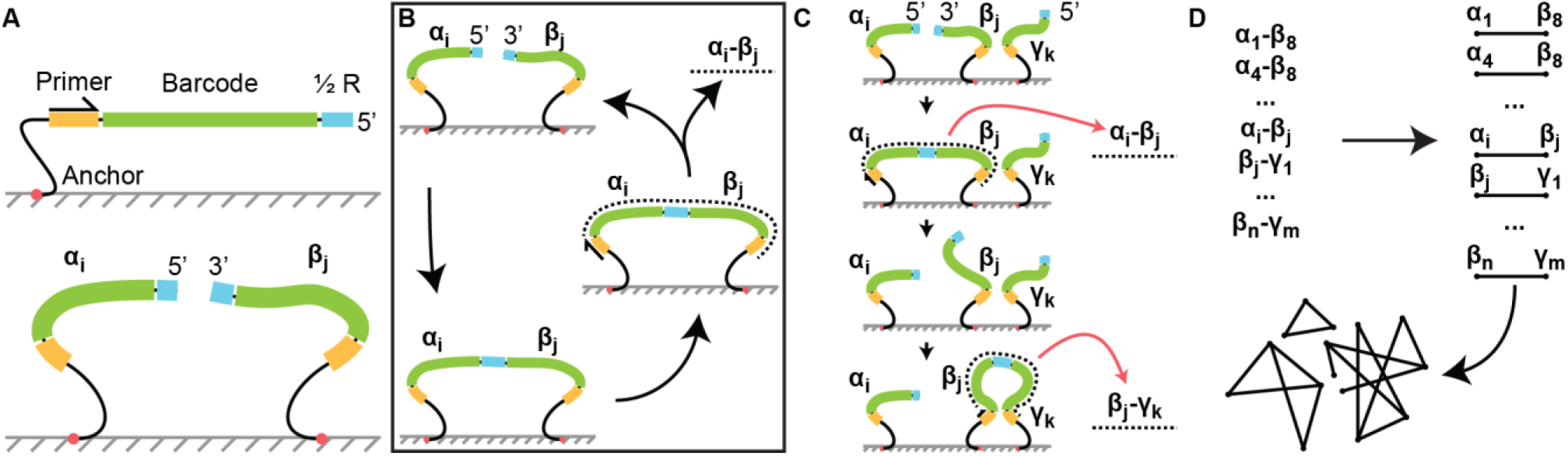
Iterative Proximity Ligation Scheme Overview. **A, top:** Each iterative proximity ligation (IPL) oligonucleotide is a single-stranded DNA synthesized to contain four components: a functional group for attachment, a primer site, a unique barcode sequence, and a restriction half-site. **A, bottom:** The oligonucleotide population is an approx. 1:1 mixture of two opposite polarities (**A, bottom**) such that they can be ligated using *e.g*. T4 ligase in the presence of a short bridging oligo (not shown). Primer and restriction site sequences are shared across each polarity; barcodes are unique across the entire population. **(B)** One round of IPL consists of ligating two oligonucleotides of opposite polarity, extending one of the primers to duplicate the barcode pair into the solution, and restriction to revert both oligonucleotides to their un-ligated state. **(C)** Presence of multiple neighboring oligonucleotides allows a different barcode pair to be recorded each round. IPL acts on a bulk population of oligonucleotides in parallel, with each round recording many barcode pairs at once. **(D)** After multiple IPL rounds, all recorded barcode pairs are sequenced by next-generation DNA sequencing. Interpretation of sequenced pairs as a graph – with each individual barcode treated as a node and each pair treated as an edge – is a rich source of information about the barcoded neighborhood.

We first experimentally demonstrate proof-of-principle iterative ligation on a simple model system.

## Results & Discussion

### Initial instantiation of iterative proximity ligation (IPL)

In order to demonstrate the feasibility of iterative proximity ligation (IPL), we used streptavidin to immobilize biotinylated oligonucleotides adjacent to one another. The oligonucleotides were designed such that their ligation could be reversed by the addition of a restriction endonuclease, leading to potential rounds of ligation, cleavage, and re-ligation (**Table 1**). We performed one such round, using qPCR to quantify the proportion of ligated products after each reaction (**Fig 2**). Six replicates of these experiments were performed.

**Table 1.**
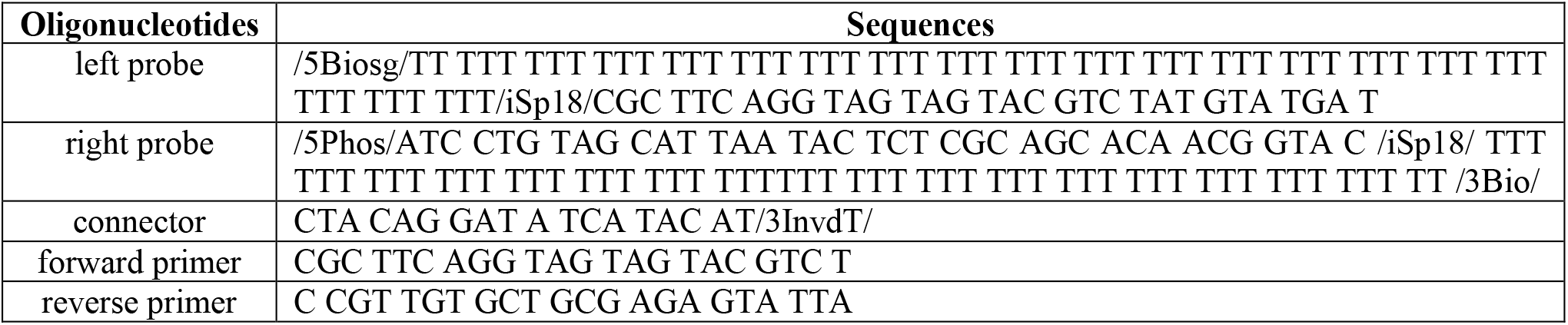
Oligonucleotides used in this work.

**Fig 2.**
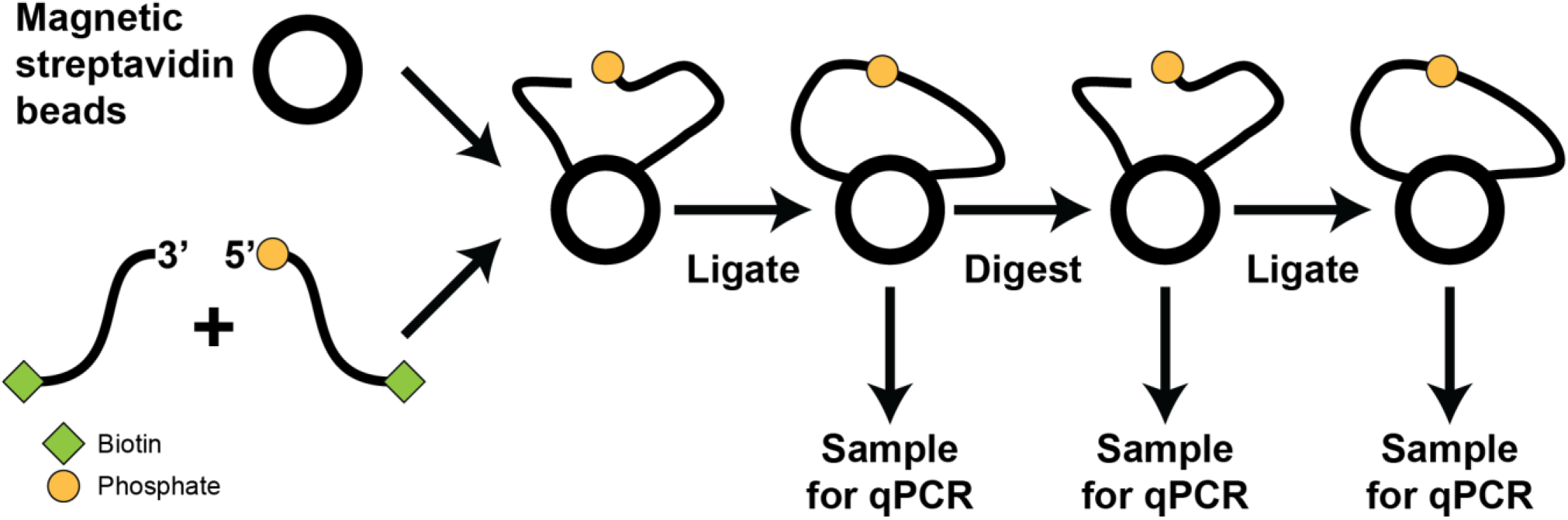
Testing reversible ligation on solid support. Left and right probes (**Table 1**) were mixed in approx. a 1:1 ratio and then incubated with magnetic streptavidin beads. After the first ligation, beads were mixed well and randomly sampled by pipette for qPCR. The remaining beads were washed and incubated with a restriction enzyme, and sampled again. A second ligation was carried out and sampled. All ligation and restriction steps required addition of a connector complementary to the restriction/ligation site (not shown; **Table 1**).

Initially, biotinylated left probe and right probe (2.5 μL of 10 μM stocks, each) were mixed and incubated with 2 μL streptavidin beads in 1x B&W buffer, with final probe and bead concentrations adjusted to ensure there were approximately equimolar numbers of probes and streptavidin binding sites (**Supplementary Methods**). Upon the addition of T4 ligase and 1 mM ATP and further incubation for 5 min at 37 °C, a 10 μL sample was taken for qPCR, which yielded an average Cq (time to signal) of 4.6 ± 1.0 (mean ± std. dev. across six replicates) (**Fig 3B**). The beads were then washed and incubated with EcoRV-HF enzyme for 2 hours at 37 °C to cleave templates that yielded amplicons, and another 10 μL sample (equivalent in concentration to the first) was taken for qPCR analysis. The average increase in Cq for the digested samples was 6.3 ± 2.0, indicating that a significant portion of the ligated probes had indeed been digested. To regenerate the template, the beads were again washed to remove the restriction endonuclease and then incubated with a more concentrated T4 ligase for 15 min at 37 °C. The final 10 μL sample showed a decrease in Cq of 7.1 ± 1.2, indicating that cleaved probes were being religated (**Fig 3A, B**). Overall, the C_q_ values went from 4.6 ± 1.0 to 10.9 ± 1.4 upon cleavage to 3.8 ± 0.31 upon religation. We confirmed that all qPCR products corresponded to the correct size as expected from our designs (**Table 1**) by 10% PAGE (**Fig 3C**).

**Fig 3.**
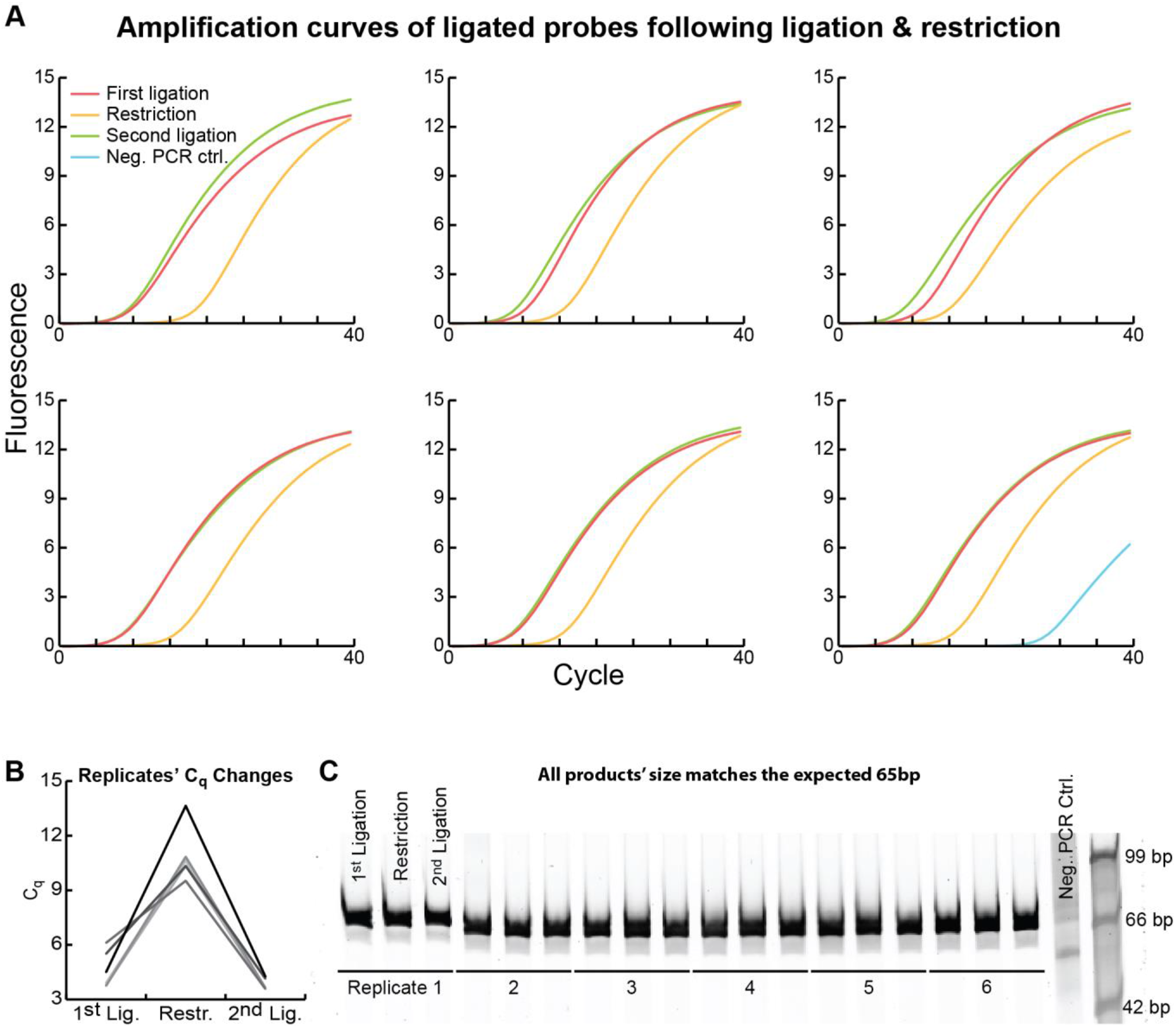
Quantitation of ligated probes following ligation & restriction. **A:** Beads with attached probes were sampled after the first ligation, restriction, and second ligation for each of six replicates. We quantified the number of ligated probes in each after each reaction using qPCR. **B:** Each replicate’s C_q_ increased significantly after restriction, implying a significant decrease in remaining ligated pairs. **C:** All qPCR reactions were run on a PAGE gel with a size marker to confirm the 65 bp product size expected from our oligonucleotide sequences (**Table 1**).

### Recovery of positional information by DNA sequencing and spring graph layouts: theoretical considerations & simulations

IPL can be used to obtain spatial information about shapes much larger than the molecules used. Here we demonstrate by simulation that we can in principle recover two-dimensional patterns by applying spring layout algorithms to IPL graphs.

First, we simulated deposition of DNA oligonucleotides on a flat glass slide as a Poisson point process^10^ within a defined geometrical shape, in this case a letter “W”, and assumed that oligonucleotides remain covalently attached only within this desired shape, with all non-specifically bound molecules washed away (**Fig. 4A**). Such shapes can, for example, be patterned across slide surfaces using photolithography^11^ or various other methods^12^. Each oligonucleotide was randomly assigned a 5’ or 3’ polarity with equal probability. For this ideal scenario, we assumed after multiple rounds of IPL that any probe pairs of complimentary polarity sufficiently close to each other to ligate did so at least once with certainty. Only the list of barcode pairs thus obtained was used to recover the layout: no additional knowledge whatsoever about oligonucleotide positions was used. We used the Graphviz^13^ Neato implementation of the Kamada-Kawai (KK) algorithm^14^ to recover the layout (**Fig. 4B**). The algorithm initializes node positions randomly and thus may cause topologically twisted layouts (**Supplementary Fig. 1**), so we pursued a layered layout strategy. We identified the graph center – defined as the set of nodes with graph eccentricity equal to graph radius^15^ – and used a randomly selected central node as a seed to layout the neighborhood of radius 1 around it. We then used the positions of nodes in this layout to initialize the layout the neighborhood of radius 2. We repeated this cycle until all nodes were laid out. The recovered shape was scaled such that the median distance between oligonucleotides in the recovered layout was equal to the median distance in the original shape. Note that due to the nature of the Poisson point process this median value is, except in pathological cases, dependent only on the IPL oligonucleotide lengths and hence highly conserved between different shapes (**Methods**). To measure large scale shape distortion in the layout versus the actual oligonucleotide positions, we aligned the original shape and the recovered layout. First, we chose the node closest to the centroid of the actual shape as a coordinate origin. The same node (red spot in **Fig. 4B**) in the layout was aligned to the coordinate origin, and we found the optimal rotation (within 0.1° resolution) that minimizes the sum of Euclidean distances between nodes’ original and recovered positions. For each node we then measured the Euclidean distance between its original and recovered positions (**Fig. 4B, heatmap**).

**Fig. 4.**
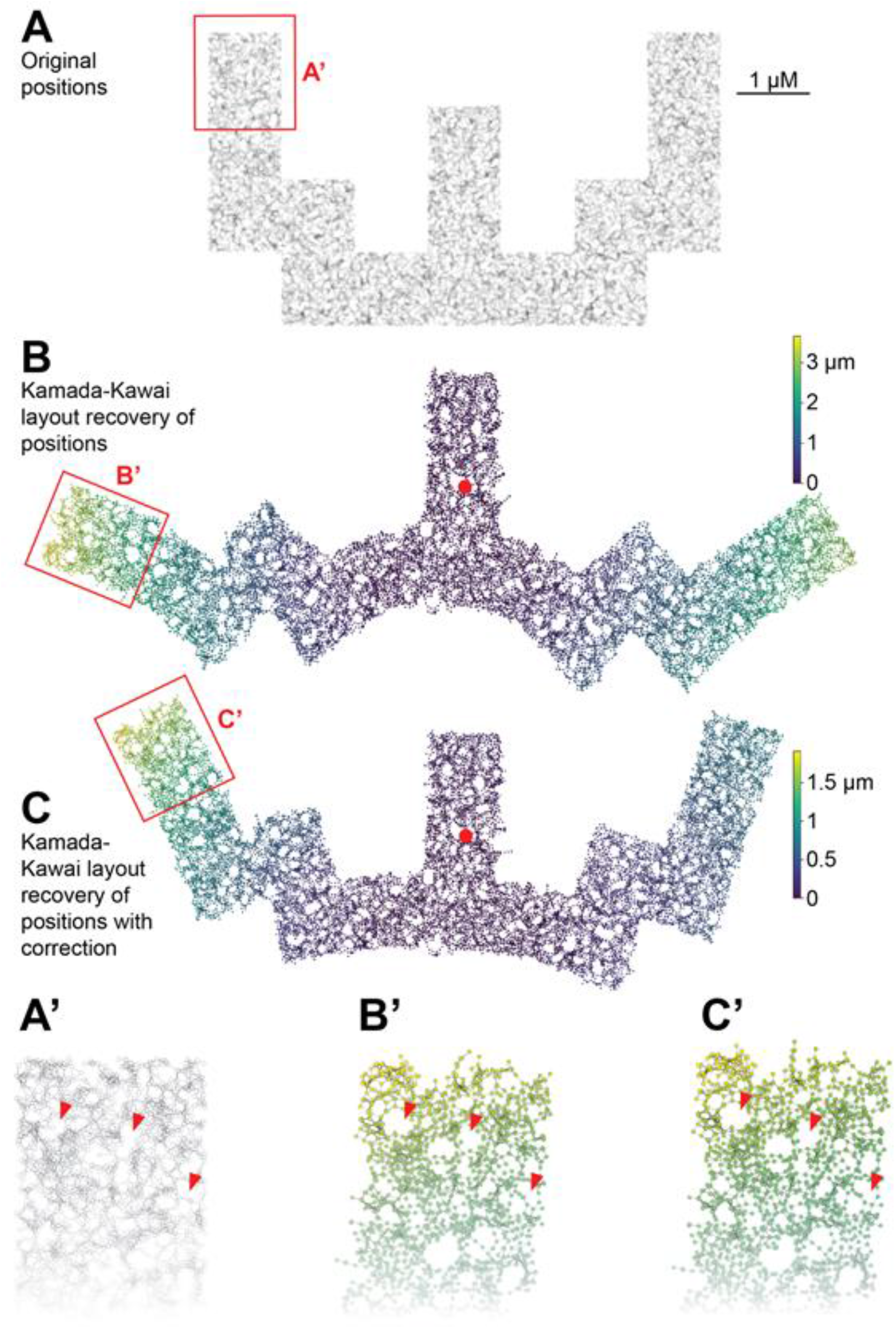
Simulations demonstrating recovery of spatial information from DNA sequences using spring-layout algorithms. **(A)** DNA as might be deposited on a surface with a photolithographed “W” shape. **(B)** Given only an IPL graph and no other positional information whatsoever, we recovered (a distorted version of) the original shape. **(C)** We can improve layout fidelity by applying to the recovered layout an additional round of a localized variant of the Kamada-Kawai algorithm. **(A’-C’)** Local topology is recovered, albeit with some distortion.

The primary cause of distortion is the discrepancy between the fundamental assumptions of the KK algorithm and concave geometries (**Supplementary Fig. 2**). To correct for this, we took the recovered layout and re-fed it into a modified variant of the KK algorithm that performs local-only spring energy minimization instead of a global minimization. This significantly decreased large-scale distortions (**Fig. 4C**).

Using this approach, we recovered all the letters for message “HELLO WORLD” (**Supplementary Fig. 3**). Note that the orientation of some letters is inverted and that this is equally consistent with the information provided: we have no way of resolving this ambiguity without additional information such as fiduciary markers or connections spanning the letters.

KK layout of two-dimensional patterns recovers not only large-scale structure (**Fig 4C**) but also finer structural details (**Fig 4A’-C’**). To quantify layout fidelity at smaller scales, we randomly sampled 50 nodes (oligonucleotides) from each letter in “HELLO WORLD” and examined their neighborhoods of graph radius 10, corresponding to a physical distance of approx. 419 ± 52 nm (mean ± std.dev.) from the sampled node. For each oligonucleotide in this neighborhood we compared the distance between its original and recovered position, with the sampled node acting as the common coordinate origin for the local comparison. The error in position for all nodes was 32 ± 20 nm with a median of 28 nm. We expect that nodes further on the graph from the sampled node will have a larger error: looking only at nodes 10 hops from the sampled node, the error in position was 34 ± 22 nm with a median of 30 nm.

Roughly stated, this means that recovered oligonucleotide positions within a half-micrometer radius circle deviate from their true positions by 4% of its diameter. Note that each oligonucleotide is approx. 34 nm in length, meaning that the average error in recovered position is on the order of an oligonucleotide length. As can be observed around the “holes” in **Fig 4A’-C’**, a primary cause of local distortion is the same concave geometric distortion that affects larger-scale structures, as discussed above.

### Recovery of positional information under non-ideal reaction efficiencies

The above result relies on the strong assumption that all oligonucleotides within mutual ligation radius will indeed ligate with certainty. We want to ask how robust positional recovery is when both the number of IPL rounds and ligation efficiencies are limited.

We first simulated multiple replicates of ligating a “W” shape for various numbers of IPL rounds and under various ligation efficiencies (**Fig. 5**). These simulations hint that there is a tradeoff between ligation efficiency and the number of IPL rounds needed to recover the positions of most nucleotides, and that, unsurprisingly, at least three IPL rounds are needed to be at least somewhat effective. Note that while the percent of all possible participating edges increases gradually as we use more IPL rounds and raise ligation efficiency, the number of participating nodes shows a sharp threshold effect as we transverse the parameter space. This latter effect has been explored in the context of random geometric graphs, and we refer the reader to references 16, 17, and 18 for a mathematical discussion and characterization of random graph connectivity.

**Fig 5.**
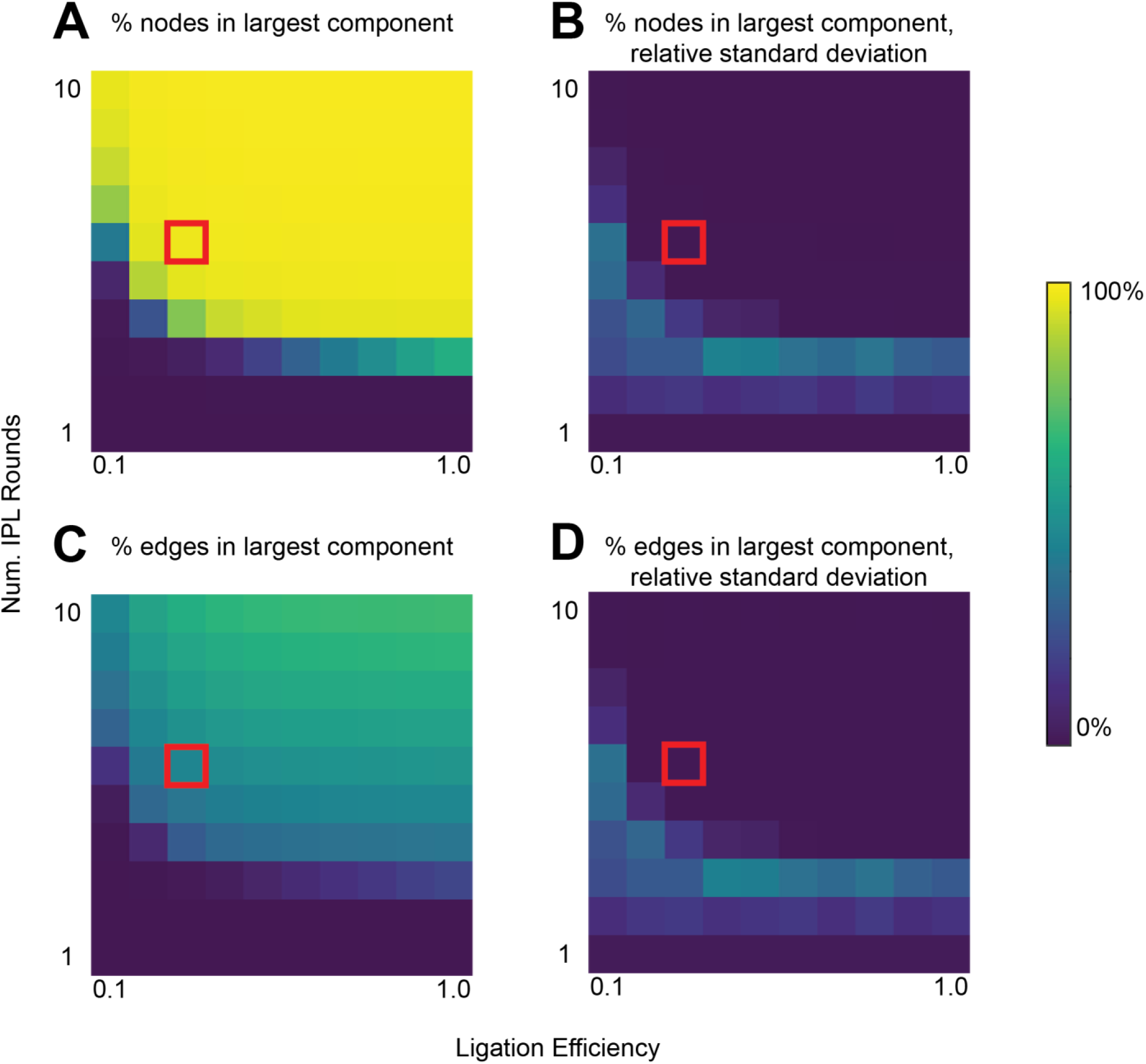
The number of possible nodes and possible edges recovered increases with the number of IPL rounds and ligation efficiency during each round. For each combination of IPL rounds and ligation efficiency parameters, we simulated 100 independent replicates of patterning a random “W” shape via a Poisson process, followed by iterative simulation of ligations between deposited oligonucleotides. We then tallied the number of nodes and edges participating in the largest IPL graph component as a percent of all possible participating nodes and edges under an ideal scenario (*i.e*. where all nodes would ligate with certainty if they were within reach of two complimentary oligonucleotides). **A:** Percent of nodes participating in the largest component, averaged across 100 simulations. **B:** Relative standard deviation across the 100 simulations in **A. C:** Percent of possible edges participating in the largest component, averaged across the 100 simulations. **D:** Relative standard deviation across the 100 simulations in C. Red square indicates parameters used for simulation in **Fig 6**, with an average of 98% of nodes and 44% of possible edges participating in the largest component.

Based on these simulations, we selected a parameter combination with a low ligation efficiency of 30% that we still expected would recover most oligonucleotide positions (**Fig. 5, red square**). Note that this appears to be below the ligation efficiency observed experimentally. We then applied the above layout approach to the resulting graph and were able to recover the shape, albeit with more distortion (**Fig. 6**). Similar results were observed for the other letters in “HELLO WORLD” (**Supp. Fig. 4**).

**Fig 6.**
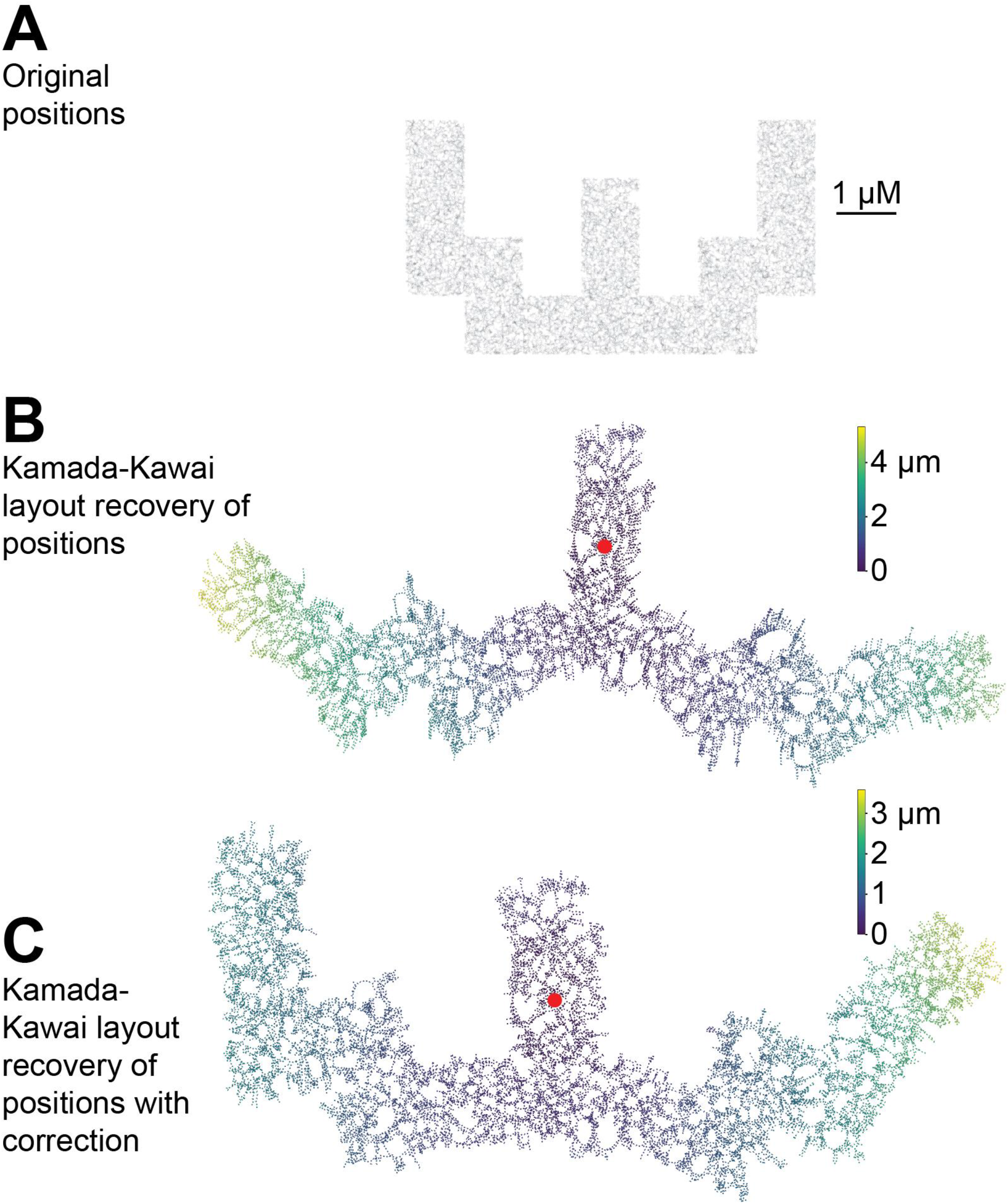
Six IPL rounds with a 0.3 ligation efficiency still recovers the original pattern overall. Note that nodes not connected to the largest graph component are not shown, as we cannot ascertain their position relative to the bulk of their connected peers.

Quantitatively comparing local distortion in neighborhoods of graph radius 10 to the ideal case above, we find that the overall error of recovered positions across all letters increased to 89 ± 56 nm with a median of 80 nm, compared with the ideal case above of 32 ± 20 nm, median 28 nm. For nodes 10 hops from the sampled nodes, the error of recovered positions increased to 113 ± 61 nm with a median of 105 nm, compared with the ideal case above of 34 ± 22 nm, median 30 nm. This means that local errors are now often larger than the reach of an oligonucleotide pair of 2 × 34 nm = 68 nm.

Based on visual inspection of the layouts, our hypothesis for a source of this distortion is that the absence of a substantial portion of possible edges creates many more “holes” in the graph without any edges that, as for the concave geometries discussed above and in **Supp. Fig. 2**, expand and push nodes – now no longer having a more direct path between them – apart. An equivalent way to think about this is that missing edges prevent KK from finding rather direct graph paths between proximal nodes, forcing it to use more circuitous ones, and this mimics the case in which the nodes are further apart in Euclidean space than they really are. This expansion hypothesis is supported if we look at how areas of local neighborhoods expand during the layout: we take from each letter the same 50 random nodes used for the local distortion metrics above, and calculate the physical area bounded by the polygon of nodes of graph distance 10 from each sample node, for both their original and recovered positions.

For the ideal case above where all possible edges were present, the ratio of areas of recovered to original bounding polygons was 1.01 ± 0.08 (mean ± std. dev.), *i.e*. the areas are conserved. For the non-ideal case, these ratios increased to 1.7 ± 0.24, *i.e*. almost doubling in area. This is visually apparent in **Fig. 6** and **Supp. Fig. 4**, where the recovered shapes are visibly larger versus the original layout despite the median distance between nodes scaled identically. In a real-world application, the practitioner will have to make a decision about how to resolve these discrepancies. One approach could be, for example, to scale the recovered shape such that the ratio of recovered areas at some scale of interest is 1:1, with the tradeoff that the distribution of edge lengths would be distorted significantly shorter.

## Conclusions

We have shown that assigning physical positions of spatially distributed molecules with barcoded DNA sequences and then recording their pairwise proximities by iterative proximity ligation can in principle reveal the relative spatial positions of the labeled objects. The underlying reason this works is because *iterative proximity ligation establishes a direct relationship between spatial position and sequence information*, by recording local molecular neighborhoods as DNA sequence pairs.

Using graph theory and simulations we have shown that oligonucleotide positions can be recovered using spring layout algorithms. Error rates in positional recovery are relatively small even with only six IPL rounds at low ligation efficiencies: approx. 1 μm diameter neighborhoods are recovered with an average oligonucleotide positional error on the order of the length of a ligated oligonucleotide pair. On scales substantially larger than individual oligonucleotides, we can recover patterned shapes. Notably, the algorithm we use for positional recovery have been implemented in three dimensions^13^. This hints at a possibility that our approach can be used to recover three-dimensional shapes using DNA sequencing. One potential application for our approach would be to extend DNA barcode strategies for the brain connectome^19^ so that we can recover not only the logical connectome but also its spatial distribution.

## Methods

### Iterative proximity ligation on magnetic streptavidin beads

#### Materials and bead preparation

2X binding and washing (B&W) buffer was prepared per Invitrogen as 10 mM Tris-HCl (pH 7.5), 1 mM EDTA, 2 M NaCl.

Biotinylated left and right probes (10 μM; Table 1) were mixed in 1x B&W buffer in a total volume of 30μL to concentrations of 0.78 μM each. The mixture was incubated with 2μL of stock Invitrogen Dynabeads M-270 Streptavidin (Invitrogen catalog #65305) at room temperature for 20 minutes to allow immobilization.

After binding, beads were washed with 200 μL of 1X B&W buffer and resuspended in 200μL nuclease-free H_2_O.

#### Iterative proximity ligation

Some 22 μL of resuspended beads was combined with 22 μL of 100 nM connector (Table 1). Ligation was carried out by adding 5 μL 10x ligation buffer (NEB B0202S) and 1 μL of T4 ligase (NEB M0202S) diluted in 1X ligation buffer to 40U/μL, and incubating for 5 minutes at 37 °C.

After ligation, beads were washed using 200 μL of 1X B&W buffer and resuspended in 50 μL of nuclease-free H_2_O. 10 μL of resuspended beads were aliquoted for qPCR and stored at 4 °C.

The remaining 40 μL of resuspended beads was subjected to digestion. 1 μL of 10 μM connector, 5 μL 10X CutSmart Buffer (NEB B2704S), and 4 μL EcoRV-HF was added to the beads and incubated at 37 °C for 2 hours.

Reactions were washed using 200 μL of 1X B&W buffer and resuspended in 40 μL of nuclease-free H_2_O. 10 μL of resuspended beads were aliquoted for qPCR and stored at 4 °C.

Ligation was repeated on the remaining 30 μL of beads as above, except using 14 μL of 100 nM connector, 1 μL of 400 U/μL ligase, and incubating at 37 °C for 15 minutes.

After ligation, beads were washed using 200 μL of 1X B&W buffer and resuspended in 30 μL of nuclease-free H_2_O. 10 μL of resuspended beads were aliquoted for qPCR and stored at 4 °C.

#### qPCR from magnetic beads

qPCR of all aliquots was performed simultaneously. 3 μL from each 10 μL aliquot was combined with 3.6 μL of nuclease-free H_2_O, 1.2 μL of forward and reverse primers (each, 10 μM; Table 1), 1 μL Evagreen dye (Biotium #31000), and 10 μL 2X FastStart DNA Probes Essential Master Mix (Roche #06 402 682 001).

qPCR started with an initial enzyme activation step of 95 °C for 600 sec; followed by 40 cycles of 95 °C for 10 sec melting, 62 °C for 1 sec annealing, and 72 °C for 1 sec extension; three final steps to obtain Evagreen melting curves were at 95 °C for 10 sec, 60 °C for 60 sec, and 97 °C for 1 sec.

#### Gel electrophoresis of qPCR reactions

A 10% polyacrylamide gel was made by combining 12.5 mL 20% acrylamide (in 7 M urea), 12.5 mL TBE dilution buffer (in 7 M urea), 100 μL 10% APS, and 25 μL TEMED. 20 μL of each qPCR reaction was mixed thoroughly with 20 μL 2X loading dye (95% formamide, 10 mM EDTA, 0.025% BB) and denatured at 95 °C for 5 minutes. After denaturation, the temperature was ramped down to 25 °C at the rate of 0.1 °C/s. The denatured product was loaded into the gel and run at the voltage of 400 V for 2 h. After running, the gel was stained in 10 μL of 10000X SybrGold (Invitrogen, S11494) diluted with 100 mL H_2_O. We re-used the oligonucleotide fragments from Bhadra and Ellington^20^ as the DNA ladder.

### Simulating recovery of spatial information from DNA sequences using spring-layout algorithms, ideal case

We simulated depositing 34nm long (~100 base) IPL oligonucleotides onto letter shape patterns on a flat surface as a Poisson point process with intensity parameter λ = 5 oligonucleotides / (68nm X 68nm). Each oligonucleotide was randomly assigned 3’ or 5’ polarity. Under ideal conditions, we assumed all oligonucleotides within 2 x 34nm = 68 nm of each other and of complimentary polarities were ligated.

Representing each IPL oligonucleotide as a node and each ligated pair as an edge, we constructed a graph using *only* the possible oligo-oligo ligations obtained above. Any nodes not belonging to the largest connected component were discarded; only the largest connected component was retained for further analysis. In practice, this meant < 1% of nodes were discarded (data not shown).

We then iteratively applied Graphviz’s^13^ implementation of the Kamada-Kawai (KK) algorithm^14^ to recover the original letter shape. We identified the graph center – defined as the set of nodes with graph eccentricity equal to graph radius^15^ – and used a randomly selected central node as a seed to layout the neighborhood of radius 1 around it. We then used the positions of nodes in this layout to initialize the layout the neighborhood of radius 2. We repeated this cycle until all nodes were laid out.

To obtain a corrected KK layout, we modified networkx’s Python implementation^21^ of the KK algorithm to ignore node-node spring interactions outside of graph radius 20. We fed the layout obtained from the final round of the iterative Graphviz KK layout into this modified function to obtain the corrected layout.

For both corrected and uncorrected variants, we scaled the node coordinates such that the median node-node Euclidean distance in the layout was equal to the (known) median oligonucleotide-oligonucleotide distance between Poisson deposited oligonucleotides.

Incidentally, note that due to the properties of a Poisson point process, this median distance between ligated oligonucleotide pairs in a sufficiently dense field will approximately equal to 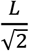, where L is the maximum possible length of an oligonucleotide pair. Here is a derivation of this claim: the median distance of ligation partners from a given oligonucleotide can be defined by the radius *r_m_* such that half of these partners lie within the circle at distances [0,*r_m_*), and half in the annulus at distances [*r_m_,L*]. The number of ligation partners within *r_m_* is, by properties of the Poisson point distribution, 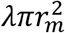 and, likewise, the number of ligation partners between *r_m_* and *L* is 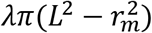. Setting 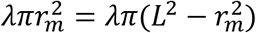 and solving for *r_m_* confirms that 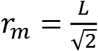. This is a useful property for scaling IPL graphs in general. Our simulations’ median distances empirically confirm this rule under various ligation regimes, albeit with a slight systematic under-estimation of *r_m_* by less than 5% (data not shown). We hypothesize this systematic under-estimation is due to edge effects: nodes near the edge of a shape have less neighbors than assumed under a Poisson distribution.

### Simulating number of edges and nodes participating in the largest component for non-ideal cases

We repeated each simulation as described below 100 times for each combination of number of IPL rounds (1 through 10) and ligation efficiency (0.1 through 1.0).

For each replicate, we simulated depositing 34nm IPL nucleotides as above in a lithographed “W” shape, with the same Poisson parameter *λ* = 5 oligonucleotides / (68nm X 68nm). As above, each oligonucleotide was randomly assigned 3’ or 5’ polarity. We then iteratively simulated the assigned number of IPL rounds. During each round, all oligonucleotide pairs within 2 x 34nm = 68 nm of each other and of complimentary polarities were iterated through in random order and each was potentially ligated at the given probability (*e.g*. ligation efficiency = 0.3 means that each ligation attempt succeeds at a 30% rate). Each oligonucleotide could at most participate in only one ligation per round.

Once all IPL rounds were simulated, we discarded all nodes not participating in the largest connected component and then tallied the percentage of original nodes and percentage of all possible edges remaining. Finally, we summarized overall node and edge participation rate in the largest connected component for a particular *IPL rounds* x *ligation efficiency* combination by evaluating the mean and standard deviation across all 100 replicates.

### Simulating recovery of spatial information (letter patterns) from DNA sequences using spring-layout algorithms, non-ideal case

We simulated re-depositing each letter pattern as in the ideal case, and then iteratively simulated IPL ligation as in the non-ideal simulations above. We then, as in the ideal case, applied the original and modified KK algorithm to the final IPL graph and again scaled the recovered positions such that the median Euclidean edge length in recovered and original layouts matched.

### Characterizing layout distortions

To characterize large scale distortion of the recovered layouts, we selected the node in the original layout closest to the centroid of all oligonucleotide positions as an anchor node. We aligned the anchor node in original and recovered layouts, and rotated the recovered layout (at 0.1° resolution) around it to minimize the sum of Euclidean distances between original and recovered node positions. To account for the recovered shape possibly being a mirror inversion of the original, we repeated this for both orientations, and selected the orientation that minimized this sum. Once the best-fitting alignment orientation and angle were chosen, we then calculated for each node the Euclidean distance between its original and recovered position.

To characterize local distortions of the recovered layouts, we randomly sampled 50 nodes from each layout and isolated the induced subgraph of neighbors within radius 10 centered around each of these nodes. We aligned the recovered layout to the original positions of nodes as for the full recovered layout above, with the sampled node playing the role of the anchor node. Once aligned, we computed the average distances between nodes’ original and recovered positions. We computed the mean, median, and standard deviation of these distances to summarize the overall distortion. We then defined *peripheral nodes* as the nodes of graph distance 10 from the sampled node. To measure distortion at a distance from each sampled node we computed the corresponding means, medians, and standard deviations for the peripheral nodes only. Finally, to compare distortions in recovered areas, we obtained the bounding polygons of the peripheral nodes and their areas for both their original and recovered positions and computed the ratios of *recovered area/original area*. These were summarized across all sampled nodes and letters by a mean and standard deviation.

### Code available on Github

Code used to produce this work and annotated demonstration Jupyter notebooks are available on our Github at https://github.com/marcottelab/mycelium.

## Acknowledgements

We would like to thank Jagannath Swaminathan and Jon Laurent for fruitful discussions and ideas. A.A.B., E.X., S.B., A.D.E., and E.M.M are co-inventors on a provisional patent covering aspects of this technology. This work was supported by a fellowship from the NSF to A.A.B. (DGE-1610403), grant 201606130011 to E.X. from the China Scholarship Council, and grants to E.M.M. from the NIH, NSF, Army, and Welch Foundation (F-1515). This work was supported by the Welch Foundation (F-1654) to A.D.E. This publication was also made possible through the support of a grant from the John Templeton Foundation (Grant 54466 to A.D.E.). The opinions expressed in this publication are those of the authors and do not necessarily reflect the views of the John Templeton Foundation.

## Supplementary Figures

**Supplementary Fig. 1.**
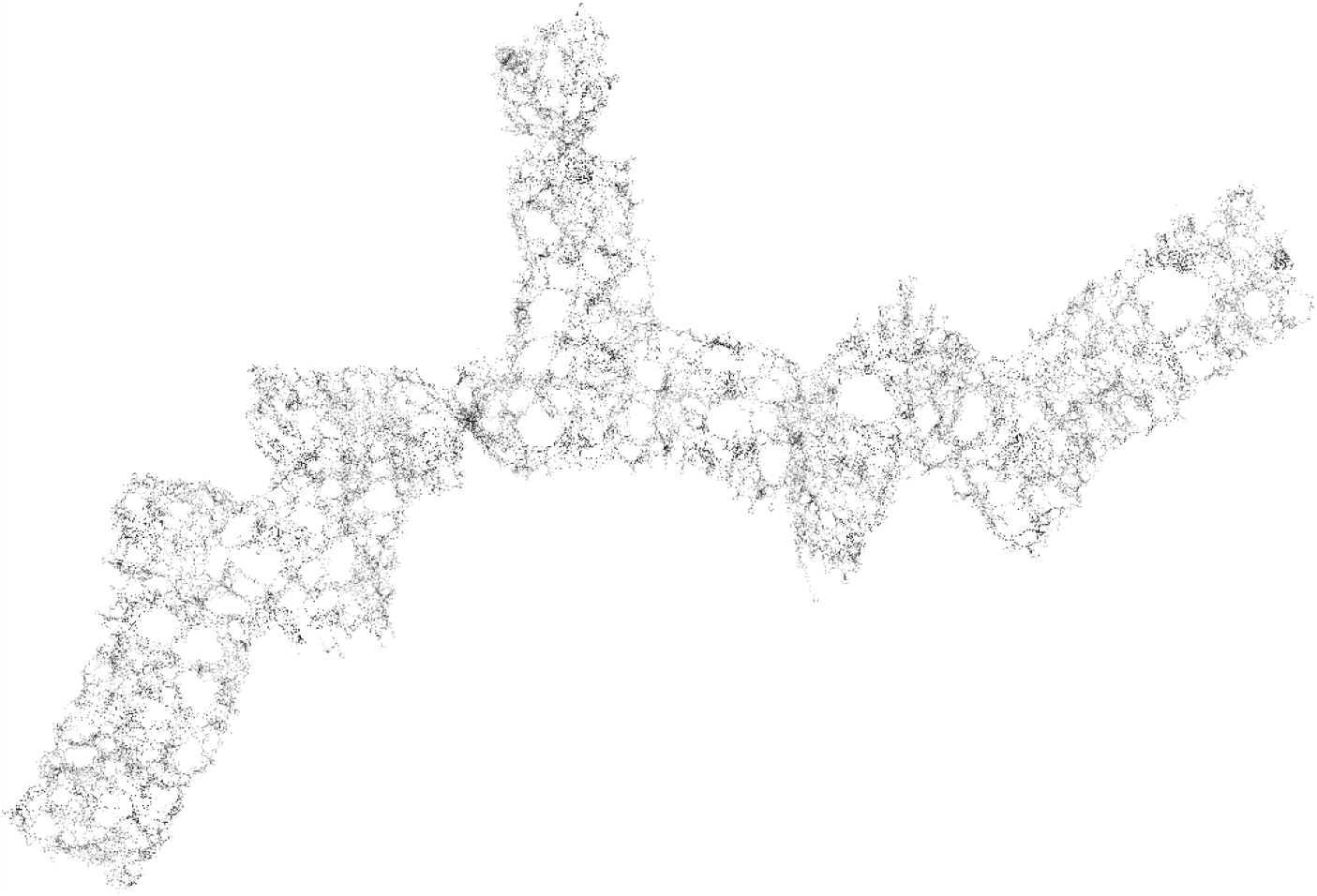
Random initialization of node positions at the beginning of a layout may cause topological twisting if nodes in various local neighborhoods of the figure happen to be initialized in opposite orientations.

**Supplementary Fig. 2.**
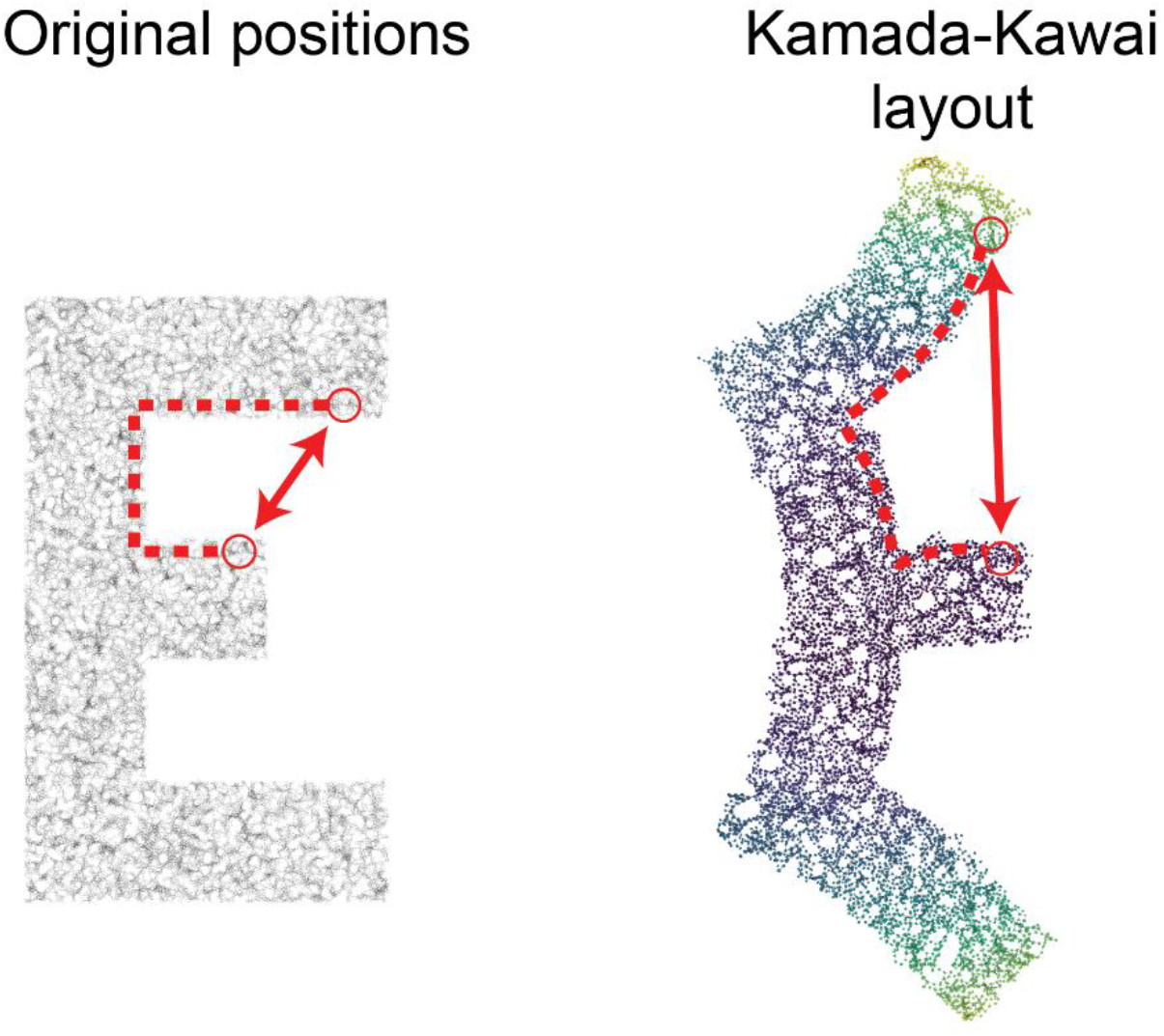
The Kamada-Kawai algorithm regards graph distance between two nodes as the desirable Euclidean distance between them. Concave geometries lead to nodes being much closer in Euclidean space (solid line with arrows) than in graph space (dashed line). This discrepancy forces nodes apart during graph layout.

**Supplementary Fig. 3.**
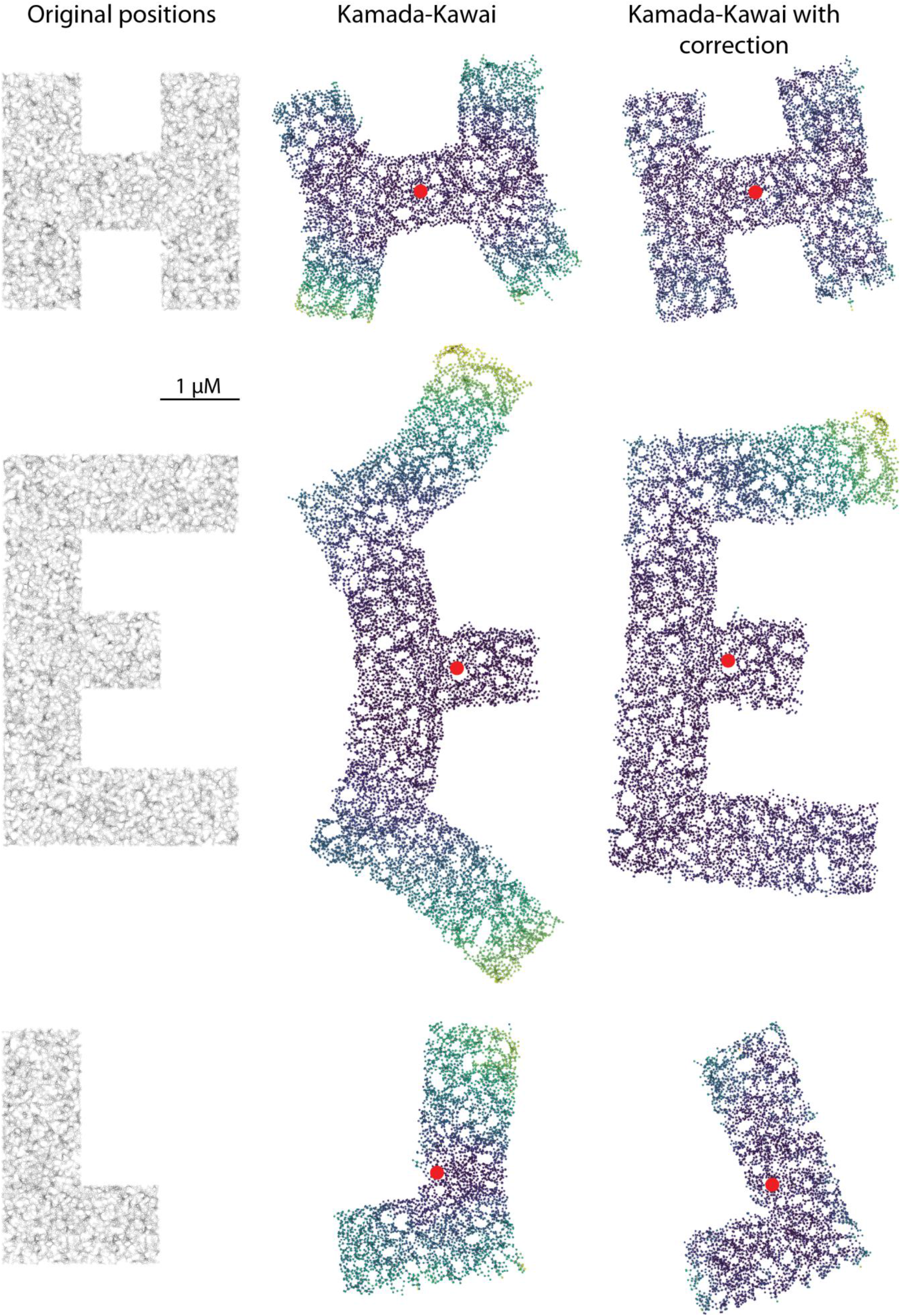
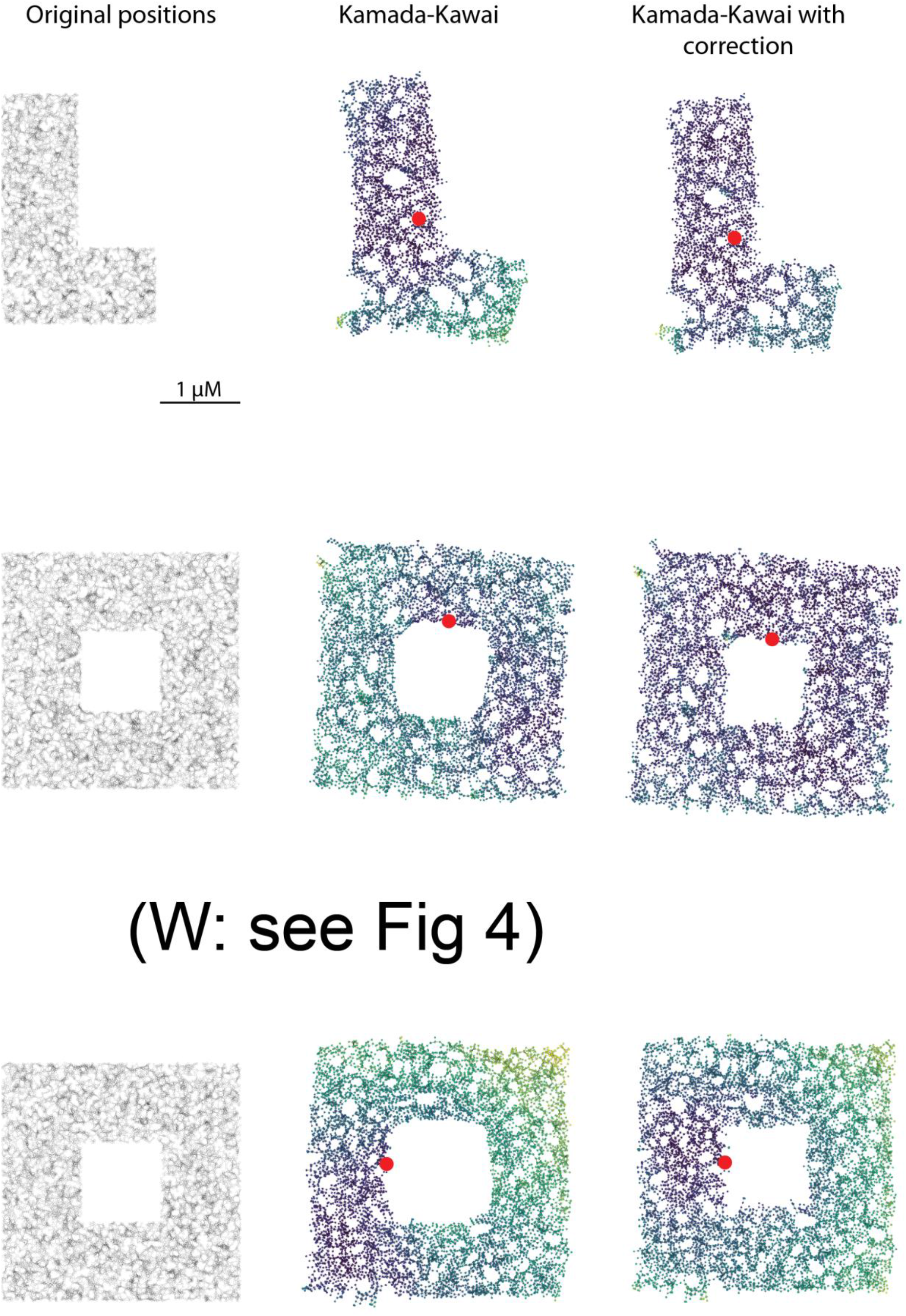
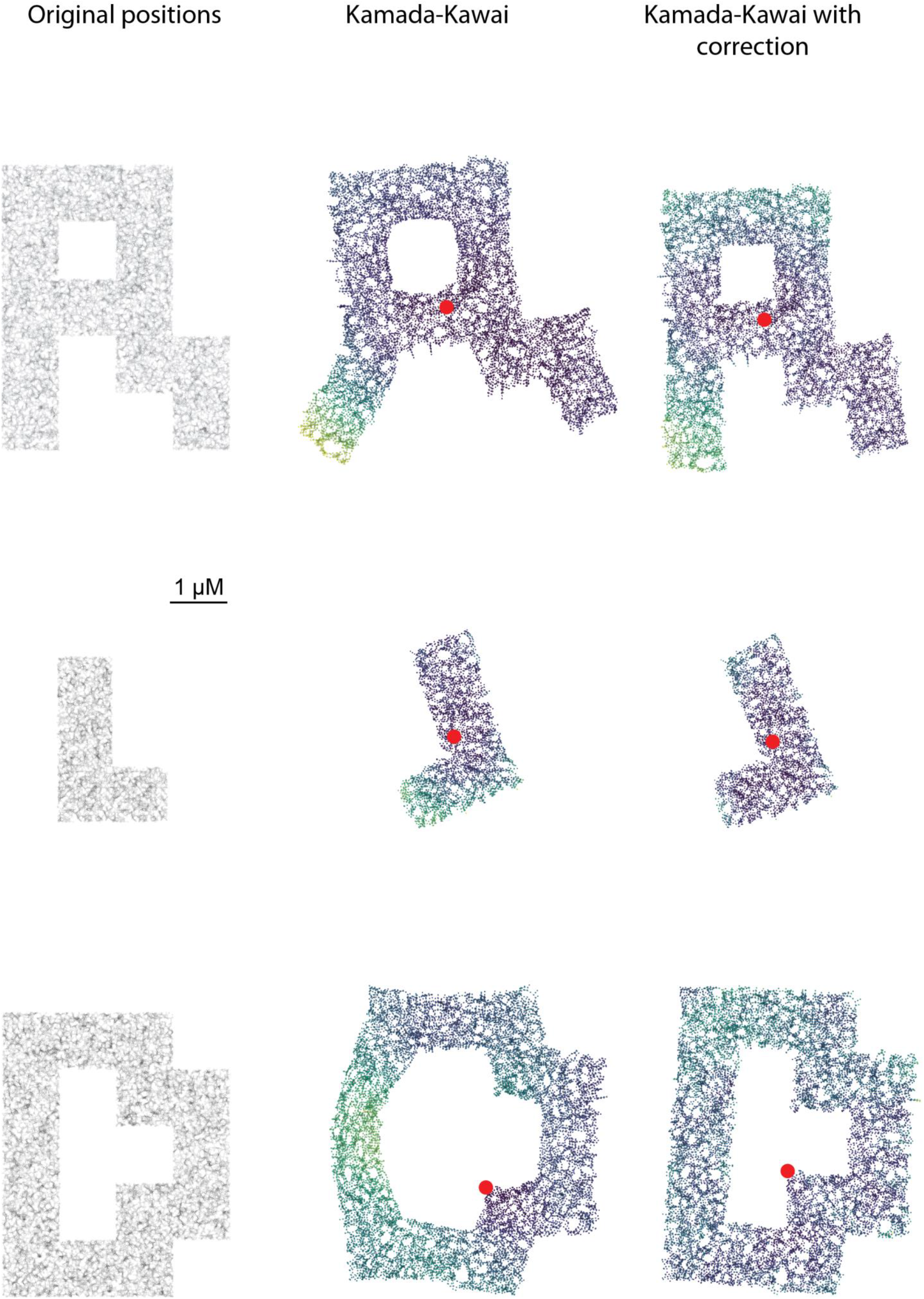

**Supplementary Fig. 4.**
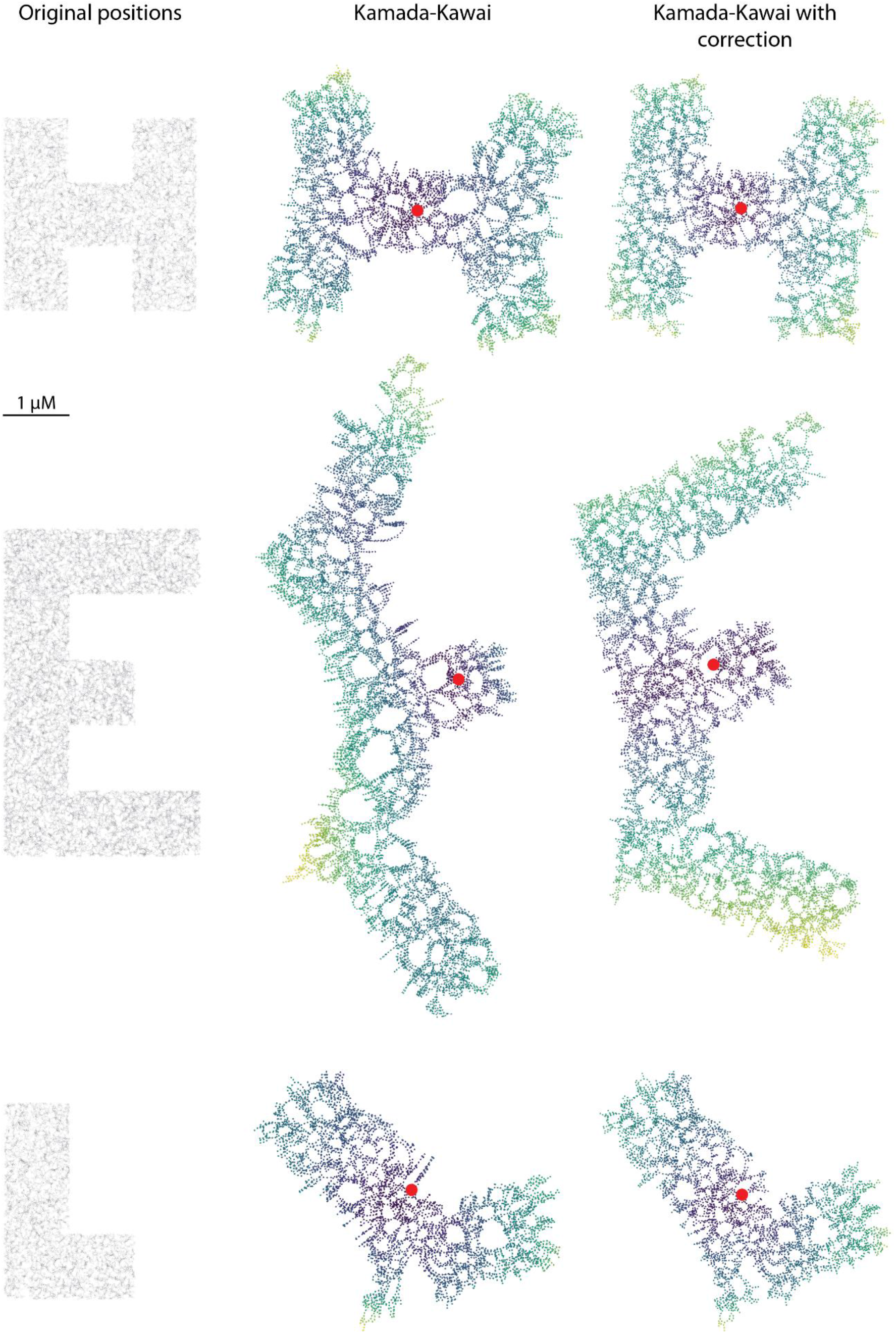
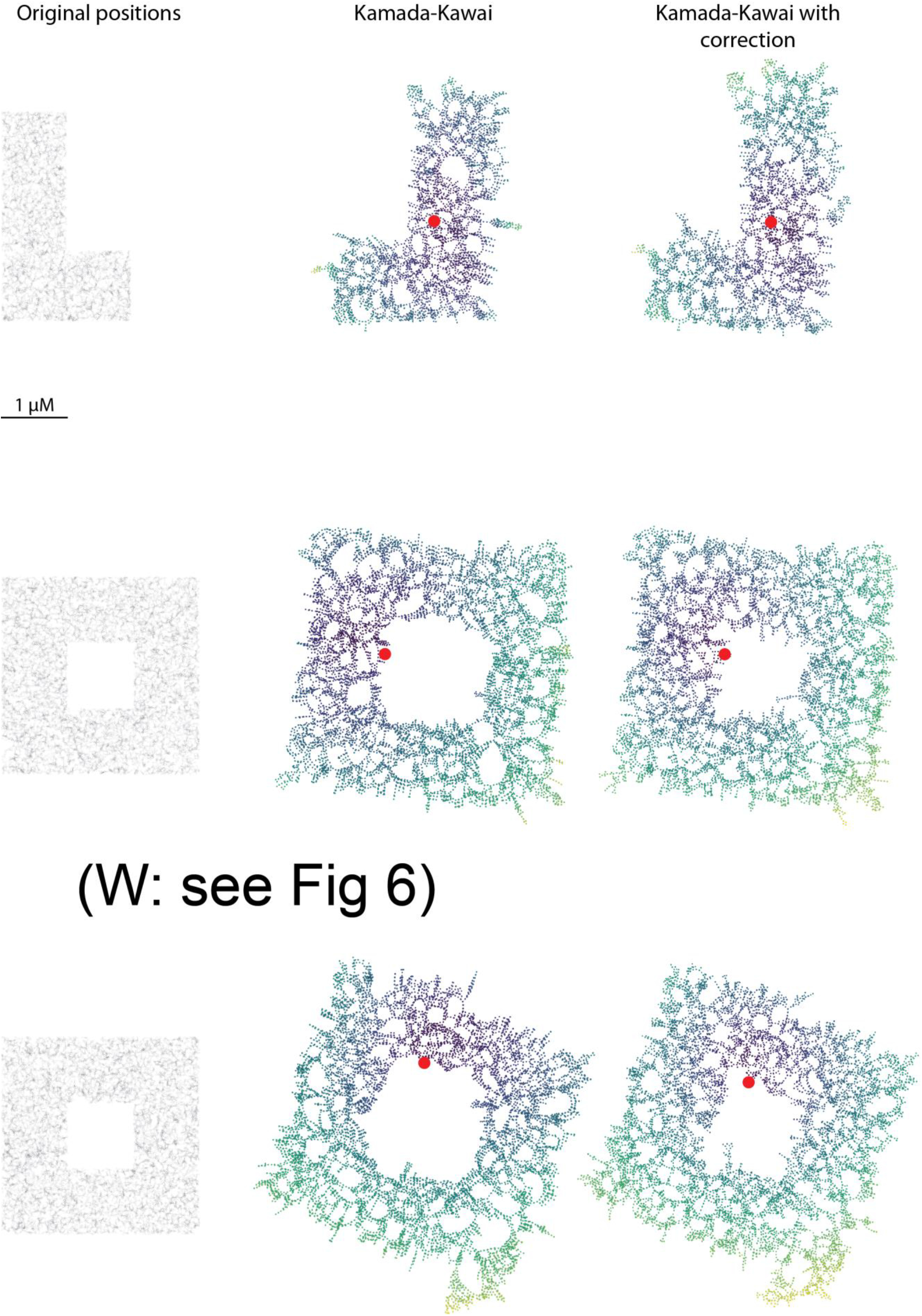
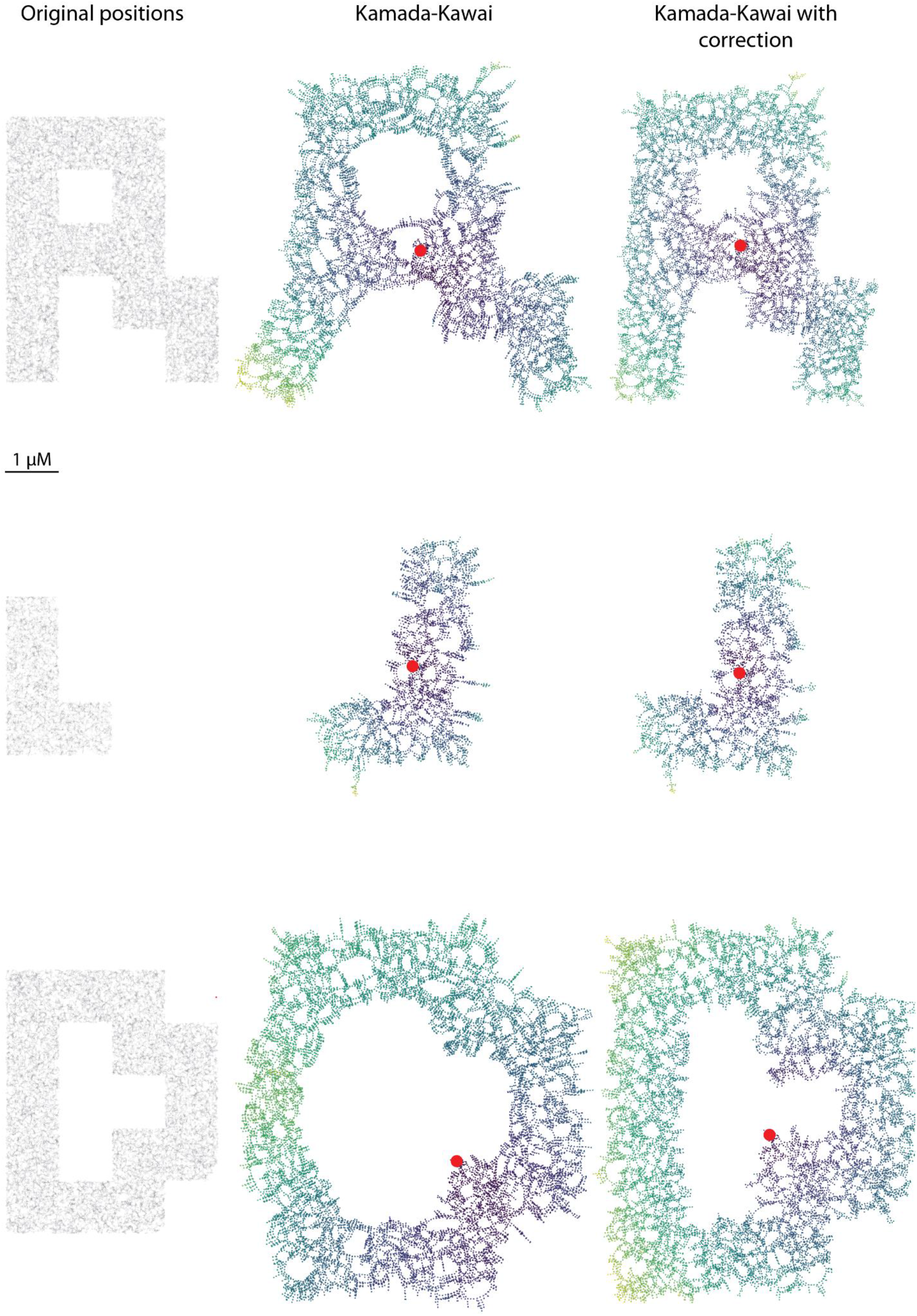

